# DELPHAI predicts heterogeneous perturbation responses with learned single-cell fitness

**DOI:** 10.64898/2026.07.01.735965

**Authors:** Xian Zhang, Hui Wu, Haikun Liu

## Abstract

Current perturbation response modelling in single-cell transcriptomics assumes conserved cell mass and loses gene expression information to latent-space decoding. We propose DELPHAI, training a fitness network and an optimal transport network jointly without biological priors, and during inference applying a fitness-gated transport with a direct gene-space retrieval. Demonstrated across two benchmark frameworks, DELPHAI ranks first in predicting differentially expressed genes, while revealing which cell lineages a perturbation depletes.

---

Tissues comprise heterogeneous cell populations that respond differently to genetic and pharmacological perturbations, underpinning essential biological functions and clinical outcomes. This complexity, captured by single-cell RNA sequencing (scRNA-seq) of control and perturbed samples, presents a challenge to *in silico* modelling, with state-of-the-art methods performing poorly [1–3]. Two conceptual limitations are shared by existing methods. First, they assume every control cell has a counterpart in the perturbed population — either mapping it onto a perturbed cell under mass-conserving optimal transport [4–6], or shifting it uniformly in latent space [7–9] — an assumption that is biologically incorrect when perturbations deplete specific lineages. Although optimal transport with relaxed marginals has been explored in single-cell genomics [10–12], no method learns cell-level survival directly from the transport objective on perturbation data. Second, the high dimensionality of scRNA-seq requires embedding into a tractable latent space, and decoding predictions back to gene expression incurs information loss that directly limits gene recovery [7, 13–15].

We therefore developed DELPHAI (Deep Explainable Perturbation Heterogeneity-Aware Inference), combining fitness-weighted optimal transport with a gene-space retriever. First, normalised gene-expression matrices are reduced to a low-dimensional embedding by principal component analysis (PCA; Fig. 1a), which performs comparably or better than non-linear embeddings such as scGen [7] and scGPT [14] (Supplementary Fig. S2). Second, a fitness network — a multi-layer perceptron mapping each control cell’s PCA embedding to a scalar weight — reweights the control distribution, and a transport network learns an optimal map from this reweighted distribution to the perturbed one; both are trained jointly against a shared Sinkhorn loss (Fig. 1b, top). Survival emerges as the loss-minimising solution: when transporting a depleted cell to a survivor incurs high transport cost, the gradient drives the fitness network to down-weight it. The resulting scores encode each control cell’s relative likelihood of surviving the perturbation, learned from distributional matching alone — without cell-type annotations, survival labels, or any external priors. Third, the transport network’s training correspondences are cached as a library of per-cell perturbation deltas in full gene-expression space (Fig. 1b, bottom). At inference, fitness gating removes predicted non-survivors using an adaptive threshold derived from the training perturbed-to-control count ratio (Methods, Eq. 2); a retriever then matches each surviving cell to its *k* nearest training controls by cosine similarity in PCA space, and a softmax-weighted sum of the retrieved deltas is added directly to that cell’s gene-expression vector. Assembling predictions in gene space bypasses the latent-decoder bottleneck that limits gene-level recovery in previous methods, while the fitness scores provide a biologically explainable readout of cellular survival. See Methods for full details.

**Fig. 1.**
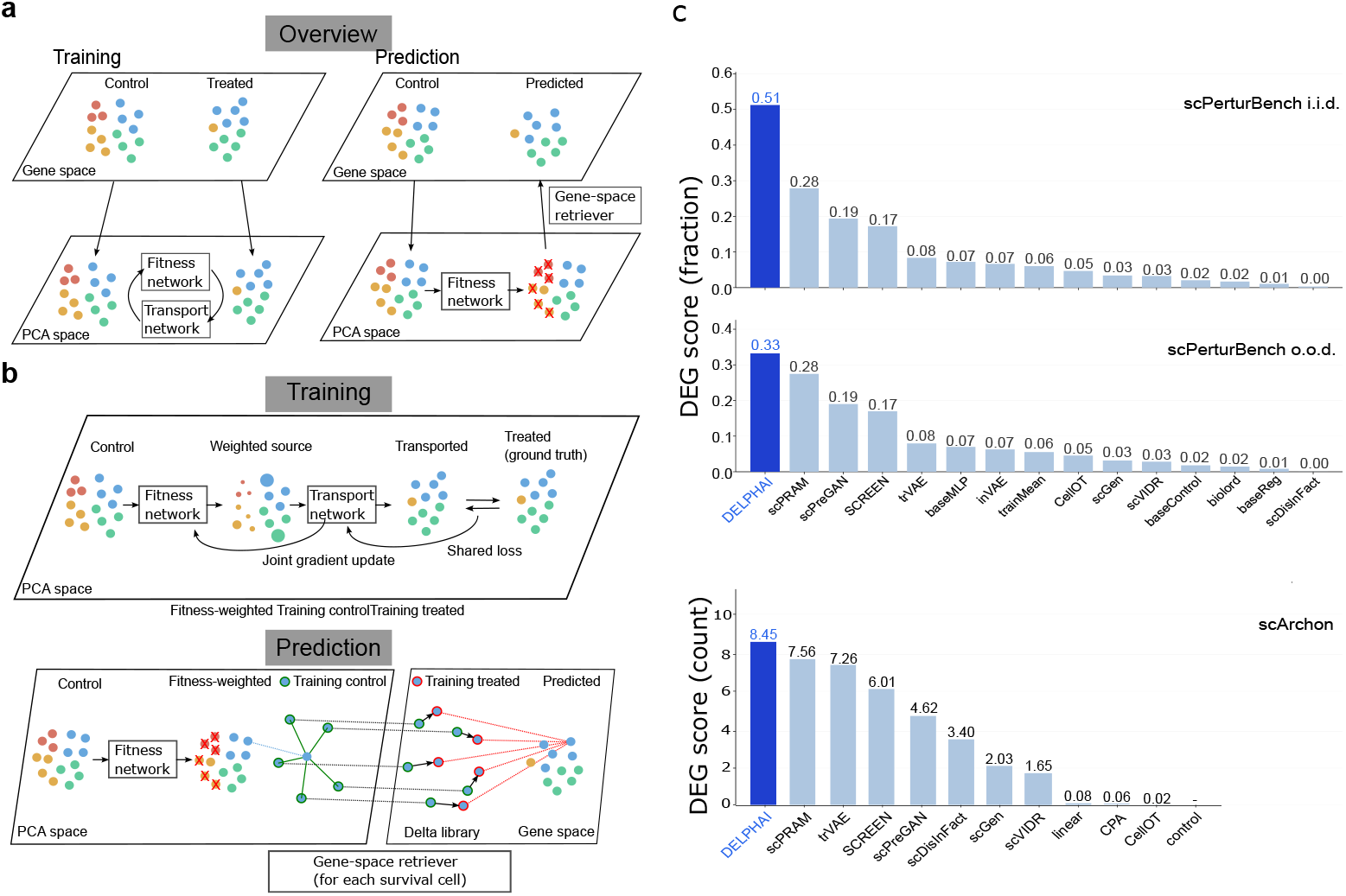
DELPHAI architecture and benchmark performance. **(a)** DELPHAI architecture overview: training (left) and prediction (right) stages. **(b)** Detailed schematics. Top: joint training of fitness and transport networks against a shared loss. Bottom: prediction — fitness gating followed by gene-space retrieval. **(c)** DEG score for DELPHAI (highlighted) against the baseline field in three benchmark settings: scPerturBench *i.i.d*. and *o.o.d*. (DEG-top-100 overlap score) and scArchon (DEG-top-20 overlap, gene count). DELPHAI ranks first in all three.

We evaluated DELPHAI in parallel on two benchmarking frameworks — scPer-urBench [2] and scArchon [3] — each with its own train/test splits, evaluation metrics, harness, and published field of baseline methods. In scPerturBench, DELPHAI is compared against 14 baseline methods on 12 datasets under two leave-one-context-out scenarios — *o.o.d*. (the cellular context held out entirely) and *i.i.d*. (half the held-out cells added back to training) — across six metrics: differentially expressed gene (DEG) overlap, Pearson correlation, mean squared error (MSE), energy distance, Wasserstein distance, and symmetric KL divergence. In scArchon, DELPHAI is compared against 11 baseline methods on six datasets under leave-one-out splits, across five metrics: top-20 DEG overlap, enrichment-term overlap, a *t*-test score, Wasserstein distance, and MSE. On DEG recovery — the most biologically informative metric — DELPHAI ranks first in all three settings (Fig. 1c): recovering 51.3% of the top-100 DEGs on scPerturBench *i.i.d*. and 33.2% on *o.o.d*., and 8.45 of the top-20 DEGs (42.3%) on scArchon — gains of 84%, 21%, and 12% over scPRAM [16], the consistent runner-up in each. Aggregated across all metrics, DELPHAI ranks third, first, and second, respectively (Supplementary Tables S1, S2, and S3).

To examine DELPHAI’s performance and explainability in detail, we focus on the clinically relevant *o.o.d*. held-out-patient setting and select two contrasting perturbations from scPerturBench: the cytokine IFN-*β* on KangCrossPatient, which stimulates immune cells without cell death, and the proteasome inhibitor CEP18770 on KaggleCrossPatient, which depletes specific cell types through cytotoxicity. Each dataset is analysed independently, holding out one patient and training on the remainder. The first held-out patient per dataset (Pat101, Pat0) is shown in Fig. 2; two further patients per dataset are shown in Supplementary Figs. S3 and S4 and support the same conclusions.

**Fig. 2.**
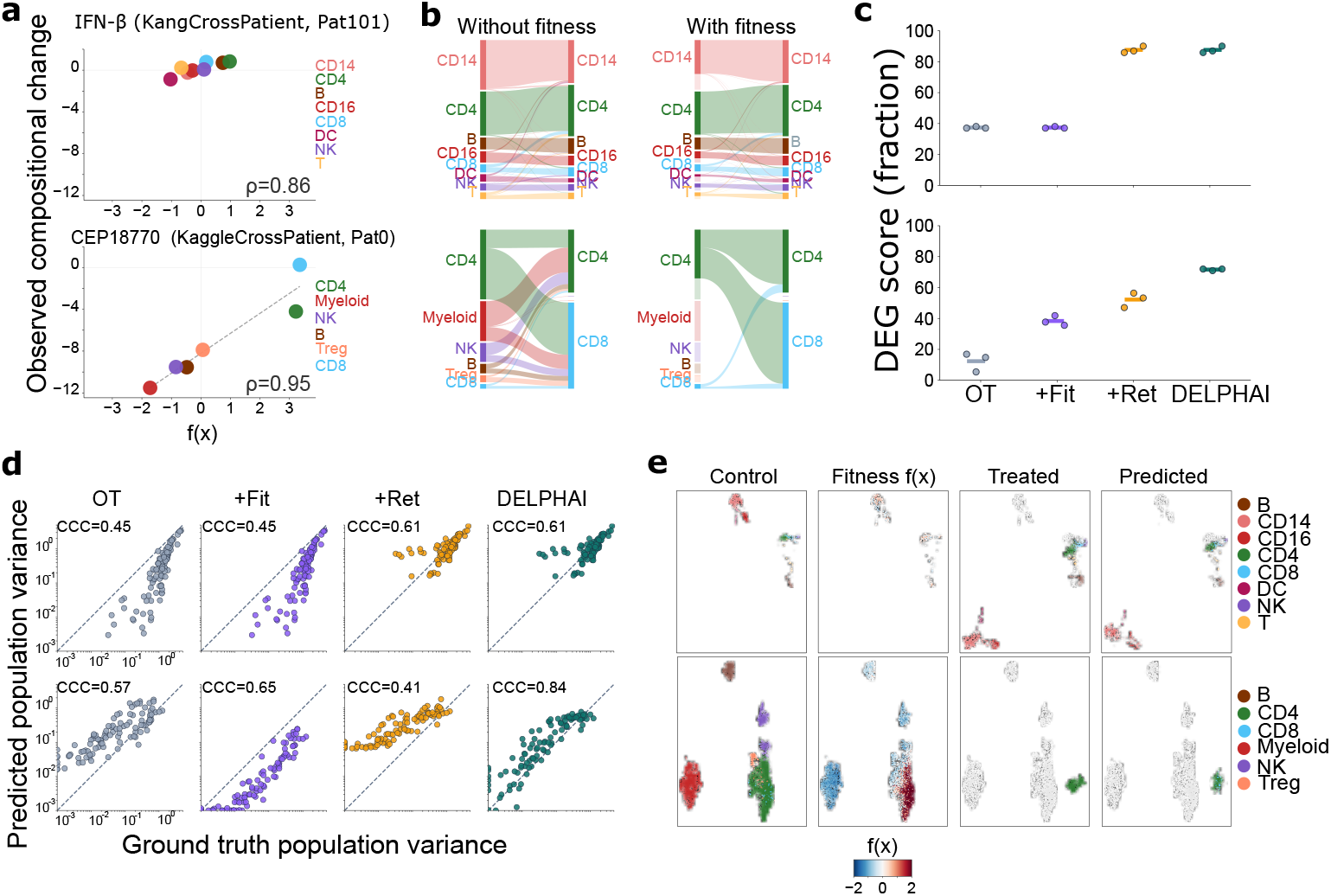
Dissecting DELPHAI’s components and predictions on two contrasting perturbations in held-out patients. All panels share the same row layout: IFN-*β* (KangCrossPatient, Pat101) on top, CEP18770 (KaggleCrossPatient, Pat0) at bottom. **(a)** Per-cell-type fitness score *vs*. observed compositional change; *ρ* values indicate Pearson correlation. **(b)** Sankey plots of transport coupling between control and treated cell populations. **(c)** Component ablation on DEG score; points are three training seeds, horizontal bars their mean. **(d)** Per-gene variance calibration: predicted *vs*. ground-truth population variance, on log-transformed axes (CCC values on panels). **(e)** UMAPs of four cell populations on a shared per-row embedding. **Control, Treated**, and **Predicted** are coloured by cell type; **Fitness** *f* (*x*) shows control cells by per-cell fitness score (red = high survival, blue = low). Treated and Predicted overlay control cells in faded grey.

The two regimes are confirmed empirically: observed compositional change is near-zero across cell types under IFN-*β* but spans *−* 12 to 0 on the log_2_ scale under CEP18770 (Fig. 2a, y-axis). DELPHAI captures both — cell-type fitness scores correlate with compositional change at *ρ* = 0.86 (IFN-*β*) and 0.95 (CEP18770; Fig. 2a, x-axis). Without the fitness gate, the transport map mis-routes depleted Myeloid, NK, and Treg cells onto surviving CD4/CD8 (Fig. 2b) — a biologically impossible transition, since these lineages do not interconvert; with the gate, depleted populations are removed before transport, preserving cell-type identity. Under IFN-*β*, gating is essentially inert.

The ablation follows accordingly. Under IFN-*β*, fitness adds no DEG benefit (0.373*/*0.373*/*0.857*/*0.857 for OT/+fit/+ret/full, means of three seeds; OT and OT+fitness are identical on every seed); gene-space assembly alone delivers the entire gain. Under CEP18770, fitness contributes at both stages — +0.250 over OT alone (0.117*→* 0.367) and +0.187 over OT+retriever (0.500 *→* 0.687; Fig. 2c). Per-gene variance calibration exposes the mechanism (Fig. 2d): under CEP18770, fitness alone under-predicts variance (CCC = 0.65; PCA reconstruction loss) and the retriever alone over-predicts it (CCC = 0.41; pooling dying and surviving cells), whereas the gated retriever is well calibrated (CCC = 0.84). Under IFN-*β* the retriever alone suffices (0.45 *→* 0.61, with no further gain from gating). UMAPs (Fig. 2e) confirm at single-cell resolution that fitness scores mark the cell types subsequently depleted, and that DELPHAI predicts both compositional and transcriptional change.

All of this emerges from control and treated populations alone, without cell-type labels or cell-mortality priors — labels are used only post-hoc, for visualisation. A single architecture recovers two qualitatively different regimes: a near-uniform fitness landscape under IFN-*β* stimulation, and a cell-type-specific survival pattern under CEP18770 cytotoxicity. DELPHAI’s per-cell fitness is therefore directly interpretable as a readout of perturbation sensitivity.

Together, DELPHAI improves the accuracy of predicted differentially expressed genes, and reveals which cell lineages a perturbation depletes; both outputs enable downstream mechanistic investigation of perturbation effects.

## Methods

### Data and preprocessing

We evaluated DELPHAI under two benchmarking frameworks. In scPerturBench [2], we used the 12 datasets of its cellular-context generalisation category, adopting its preprocessing, train/test splits, and *i.i.d*. and *o.o.d*. scenario definitions without modification. In scArchon [3], we used its six datasets (Kang, H.poly, Nault, cross-species, glioblastoma, and IFN-*α*), adopting its independent preprocessing, highly-variable-gene selection, and leave-one-out splits. Five of the scArchon datasets share source data with scPerturBench datasets and one (IFN-*α*) is unique to scArchon; because each framework applies its own preprocessing and splits, the two evaluations remain methodologically independent even where the underlying data overlap.

### Evaluation, ranking, and baselines

Baseline values are taken from each framework’s published results; no baseline was re-run or re-scored. In scPerturBench [2], DELPHAI is evaluated on the splits shared by all 14 baseline methods and scored on six metrics — PCC, MSE, E-distance, Wasserstein distance, symmetric KL divergence, and DEG-overlap score — computed on the top-100 differentially expressed genes via scPerturBench’s published pipeline (https://github.com/bm2-lab/scPerturBench). Aggregate values reported in the main text (e.g. DELPHAI’s 51.3% *i.i.d*. DEG score) are grand means (per-task scores averaged within each of the 12 datasets, then the 12 dataset means averaged with equal weight), and overall ranks follow scPerturBench (per-task rank across the 15 methods, averaged within each dataset, then across the 12 datasets and 6 metrics); per-task values for all 15 methods are in Supplementary Tables S1 and S2. In scArchon [3], DELPHAI is run under the unmodified scArchon harness on its six datasets with leave-one-out splits and scored on five composite metrics — top-20 DEG overlap and enrichment-term overlap (biological), and a *t*-test score, Wasserstein distance, and MSE (statistical); following scArchon’s procedure, methods are ranked per dataset on each metric and these ranks averaged, and DELPHAI is compared against scArchon’s published field of 11 methods, with per-dataset, per-metric values and the resulting ranking in Supplementary Table S3.

### Model

Cells are represented by log-normalised, HVG-filtered gene-expression vectors *x* supplied by each framework, and by PCA embeddings *z* of the top *n*_PCA_ = 200 principal components fit on the training-condition cells (Supplementary Fig. S1).

#### Fitness network f

R^200^*→* R_*>*0_. A three-layer MLP with hidden width 128 (Supplementary Fig. S1), LayerNorm after the first layer and LeakyReLU activations (slope 0.2). It outputs a log-fitness score per cell; weights follow by exponentiation and normalisation.

#### Transport network T

R^200^*→*R^200^. Parameterised as *T* (*z*) = *Δ φ*(*z*), the gradient of an Input Convex Neural Network potential *φ* [4]. Convexity is maintained by projecting the ICNN hidden-layer weights onto the non-negative orthant after each optimiser step and by a soft penalty on their negative parts.

#### Training objective

Given unpaired control and perturbed populations *μ*_ctrl_ and *ν*_pert_, both probability measures on R^200^, the two networks are trained jointly against a single loss:

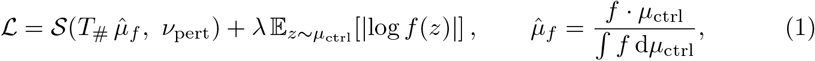

where *S* is the debiased Sinkhorn divergence (geomloss, quadratic cost, blur 0.05). Fixing the target marginal while leaving the source marginal free gives a semi-relaxed optimal transport problem, so depleted cells can be down-weighted rather than transported. The Sinkhorn term alone is invariant to a constant rescaling of *f*, since 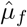 is normalised; the L1 penalty is applied to the unnormalised log *f* and therefore fixes the scale, shrinking *f* toward unity when no compositional shift is present (*λ* = 0.1, Supplementary Fig. S1). *f* is a per-cell weight relative to the population, not an absolute survival probability, and the gating (Eq. 2) uses only its relative ordering.

#### Delta library

After training, the Sinkhorn coupling *π* between the transported (surviving) training controls and the perturbed cells gives, for each training control *j*, a barycentric match 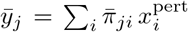 (with 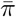 the row-normalised coupling) and hence a per-cell delta 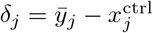 in full gene-expression space, indexed by *{z*_*j*_*}*.

### Inference

Each test control cell, with gene-expression vector *x*_te_ and PCA embedding *z*_te_, is processed in three steps.

#### Step 1 — fitness scoring

Compute *f* (*z*_te_).

#### Step 2 — adaptive gating

Let *p* be the empirical perturbed-to-control count ratio in the training data for the same perturbation, which sets the fraction of test-control cells retained. We clip *p* to [0.01, 1], so at least 1% of cells survive gating and proliferation conditions (*p >* 1) retain all of them. The gating threshold is the (1 *−p*)-quantile of *f* across the test-control population:

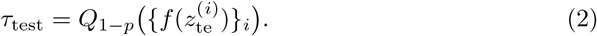

Cells with *f* (*z*_te_) *< τ*_test_ are excluded, subject to a floor of ten retained cells. The same rule applied to the training-control population yields *τ*_train_, which defines the retrieval pool for Step 3; the adaptive scheme is selected over fixed-*τ* alternatives (Supplementary Fig. S1).

#### Step 3 — retrieval and reconstruction

For each surviving test cell, let *N*_*k*_ be its

*k* nearest training controls in the retrieval pool by cosine similarity in PCA space.

Weights are computed in PCA space and applied to the gene-space deltas *δ*_*j*_:

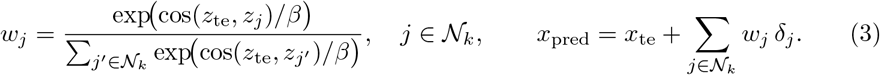

*k* and *β* are set to 5 and 0.01 (Supplementary Fig. S1). Assembling *x*_pred_ directly in gene space bypasses the latent decoder entirely.

### Ablations, hyperparameter sweeps, and statistics

#### Component ablation

A 2 *×*2 factorial design over the two DELPHAI components: OT base (neither fitness nor retriever), OT+fitness, OT+retriever, and full DEL-

PHAI (Fig. 2c). All variants share identical preprocessing, transport architecture, and training schedule. The ablation was run at three training seeds on a fixed leave-one-patient-out split; points in Fig. 2c show individual seeds and horizontal bars their mean.

#### Embedding comparison

PCA-200 shows comparable or better performance than variational autoencoder (scGen [7]) and foundation-model (scGPT [14]) embeddings on the four sweep datasets (scGPT on the two human datasets only; Supplementary Fig. S2); we adopt PCA for speed and determinism. Variational autoencoder embeddings follow the description in scPerturBench; foundation-model embeddings use the scGPT CP pretrained checkpoint.

#### Hyperparameter sweeps

Six hyperparameters are swept on four representative scPerturBench datasets (Haber, KaggleCrossPatient, TCDD, KangCrossPatient), *o.o.d*. scenario (Supplementary Fig. S1): PCA dimension *n*_PCA_ *∈ {* 10, 50, 100, 200, 300, 500 *}*; fitness-network hidden width *∈{* 64, 128, 256*}* ; retrieval neighbours *k ∈ {*1, 3, 5, 10, 20, 50 *}*; fitness shrinkage penalty *λ ∈{* 0, 0.01, 0.1, 0.5, 1.0 *}*; fixed survival quantile *τ∈ {*0.001, 0.01, 0.05, 0.1, 0.5*}*, contrasted with the adaptive-*τ* scheme; softmax temperature *β∈{*0.001, 0.01, 0.1, 1.0 . *}*

#### Variance calibration

As a complementary diagnostic in Fig. 2d, the gene-wise population variance of the perturbation delta 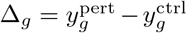 across all 5,000 HVGs is compared to ground truth via Lin’s concordance correlation coefficient applied to the log-transformed variances 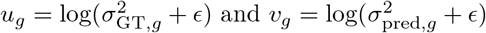 (pseudocount *ϵ* = 10^*−*3^):

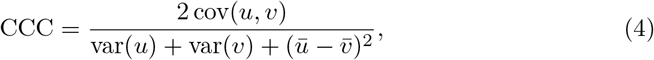

which penalises both scale bias and dispersion mismatch on the log scale.

#### Statistics and reproducibility

All cells and all perturbation tasks available in each public dataset were used; no sample-size calculation was performed. Random seeds were fixed for model initialisation and train/validation sampling. Benchmark results are single-run at a fixed seed, matching the evaluation protocol of both frameworks, which likewise fix seeds throughout [2, 3]. The component ablation (Fig. 2c) was additionally run at two further seeds to assess run-to-run variability.

## Acknowledgments

This work was supported by cloud computing credits from the AWS Activate program and the NVIDIA Inception program. We thank the authors of scPerturBench and scArchon for open-sourcing their benchmarking frameworks.

## Author contributions

X.Z., H.W., and H.L. conceived the study. X.Z. developed the DELPHAI architecture and performed the analysis. X.Z., H.W., and H.L. wrote the manuscript.

## Supplementary Information

**Table S1.**
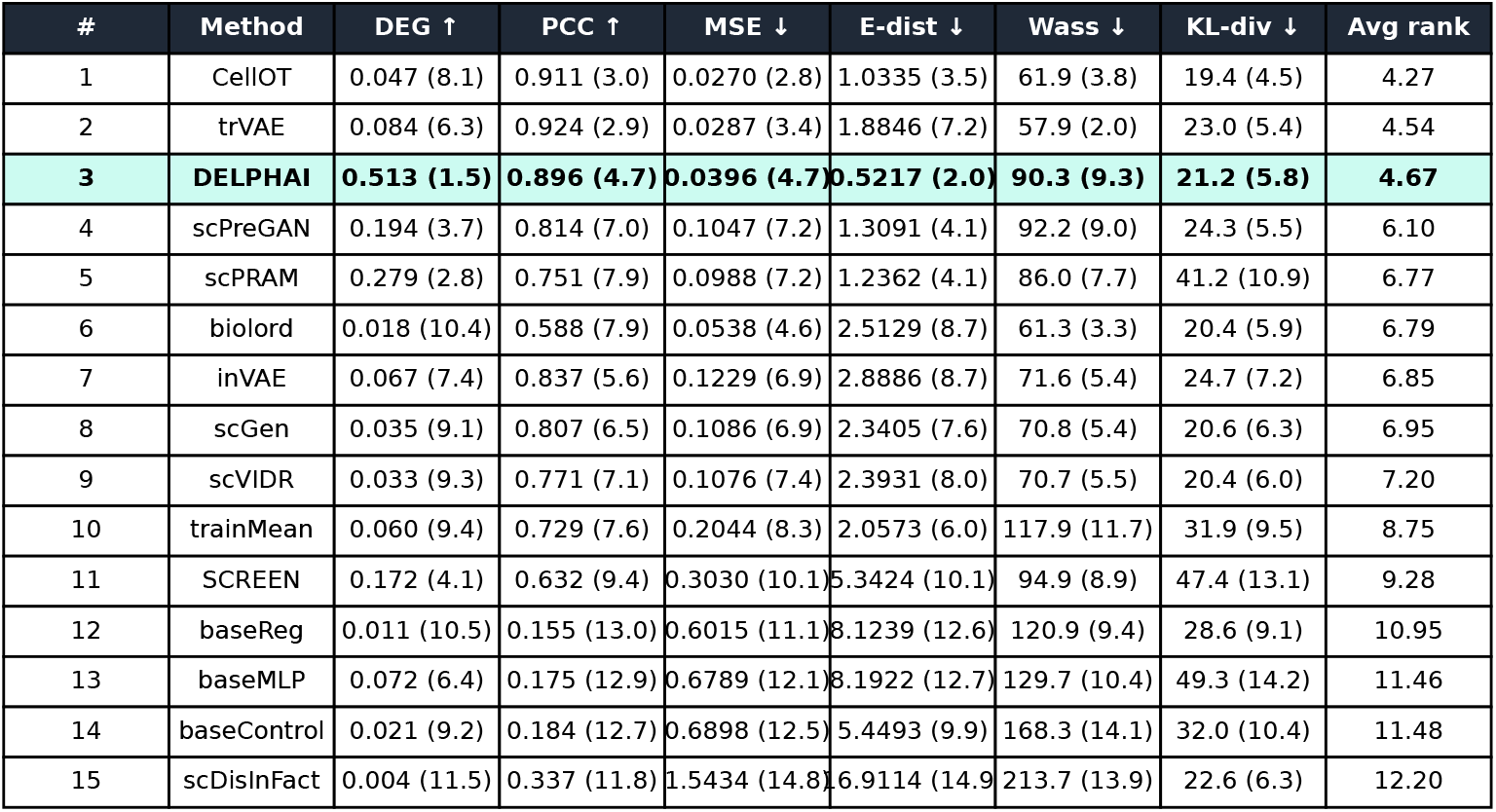
scPerturBench benchmark values, *i.i.d*.. All 15 methods ranked best-to-worst by overall mean rank. Each cell gives the metric value (grand mean across the 12 datasets) with its per-metric mean rank in parentheses; *Avg rank* is the mean of the six. Aggregation follows scPerturBench (Methods).

**Table S2.**
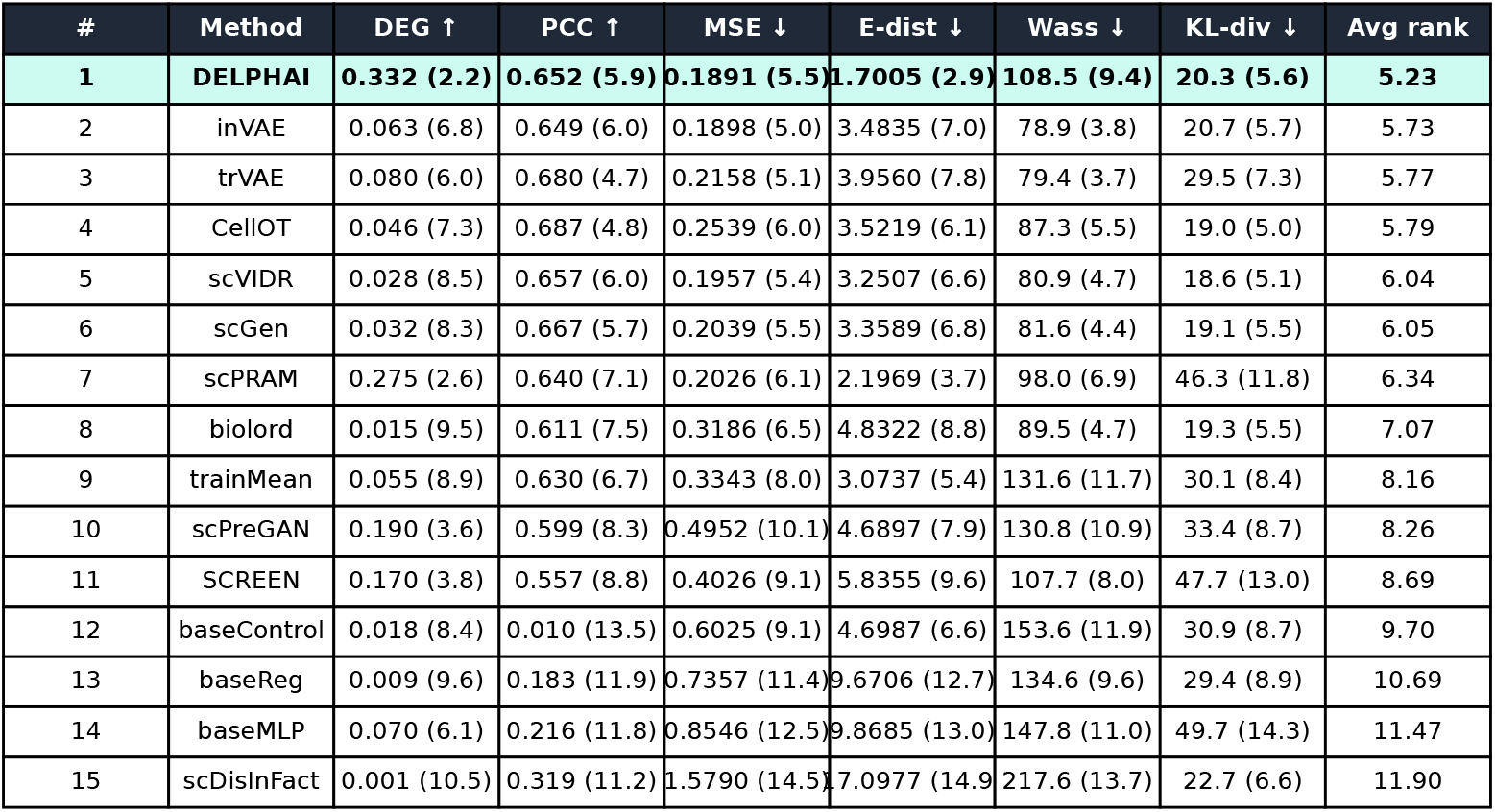
scPerturBench benchmark values, *o.o.d*. As Supplementary Table S1, but for the scPerturBench *o.o.d*. scenario.

**Table S3.** scArchon benchmark values. All 12 methods — scArchon’s nine perturbation tools plus the linear and control baselines (c2s excluded) — ranked by mean composite rank score across the six scArchon datasets [3]. Each cell gives the metric value (grand mean) with its per-metric mean rank in parentheses; ‘—’ marks metrics undefined for the control baseline. Ranking follows scArchon (Methods). DELPHAI ranks first on the biological DEG-20 overlap and second overall.

| # | Method | DEG-20 ↑ | Enrich ↑ | t-test ↓ | Wass ↓ | MSE ↓ | Overall (rank_score ↑) |
| --- | --- | --- | --- | --- | --- | --- | --- |
| 1 | trVAE | 7.26 (3.2) | 46.1 (3.6) | 7.26 (3.8) | 718 (2.8) | 0.0126 (4.3) | 0.762 |
| 2 | scPRAM | 7.56 (2.7) | 55.8 (5.4) | 6.74 (5.0) | 920 (6.3) | 0.0179 (6.2) | 0.636 |
| 3 | **DELPHAI** | **8.45 (1.8)** | **94.1 (3.2)** | **5.71 (2.5)** | **1859 (10.0)** | **0.0378 (7.0)** | **0.635** |
| 4 | scVIDR | 1.65 (6.7) | 19.7 (7.0) | 11.99 (8.0) | 743 (2.2) | 0.0134 (3.0) | 0.613 |
| 5 | scDisInFact | 3.40 (5.0) | 15.1 (7.4) | 13.89 (8.0) | 724 (2.8) | 0.0148 (5.2) | 0.586 |
| 6 | scGen | 2.03 (6.5) | 19.0 (6.8) | 12.27 (7.7) | 774 (3.2) | 0.0146 (3.8) | 0.585 |
| 7 | control | — | — | 3.89 (1.3) | 1295 (9.8) | 0.0146 (6.7) | 0.550 |
| 8 | SCREEN | 6.01 (3.5) | 65.6 (4.0) | 9.48 (7.0) | 877 (5.8) | 0.0256 (8.8) | 0.550 |
| 9 | CPA | 0.06 (8.8) | 2.2 (7.6) | 6.46 (4.5) | 792 (4.8) | 0.0149 (7.2) | 0.495 |
| 10 | scPreGAN | 4.62 (5.2) | 44.8 (3.8) | 12.82 (7.3) | 1675 (9.3) | 0.0471 (10.2) | 0.423 |
| 11 | linear | 0.08 (8.8) | 38.1 (5.0) | 121.55 (11.7) | 1248 (8.8) | 0.0102 (3.7) | 0.391 |
| 12 | CellOT | 0.02 (8.8) | 0.0 (10.0) | 47.23 (11.2) | 44359 (12.0) | 3.3138 (12.0) | 0.105 |

**Fig. S1.**
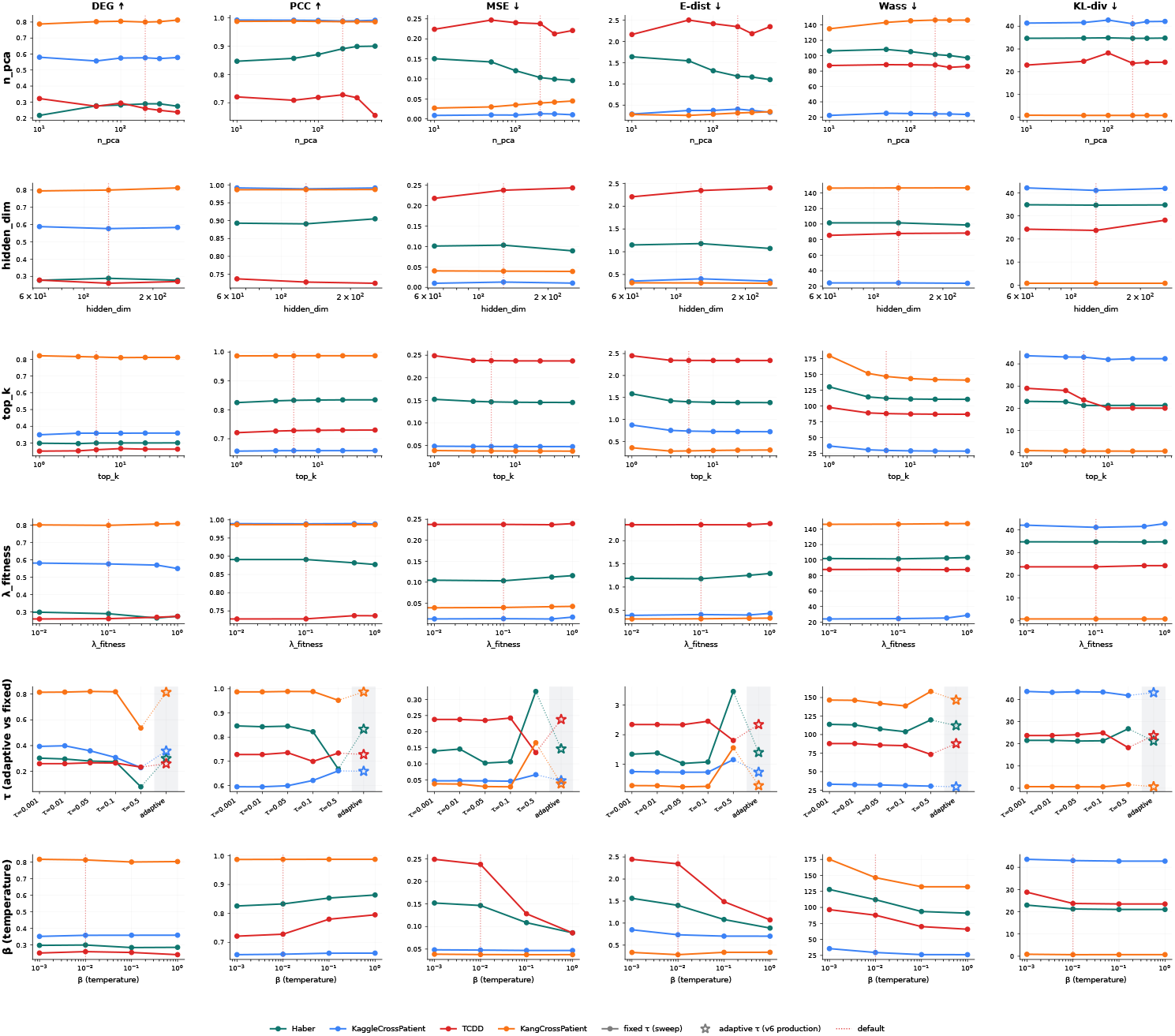
Hyperparameter sensitivity. DELPHAI’s six metrics as a function of six hyperparameters (rows: PCA dimension *n*_PCA_, fitness-network hidden width, retrieval neighbours top-*k*, fitness shrinkage penalty *λ*_fitness_, survival quantile *τ*, softmax temperature *β*), measured on four representative sweep datasets (Haber, KaggleCrossPatient, TCDD, KangCrossPatient), *o.o.d*. scenario. Lines show the per-parameter-value mean across each sweep dataset.

**Fig. S2.**
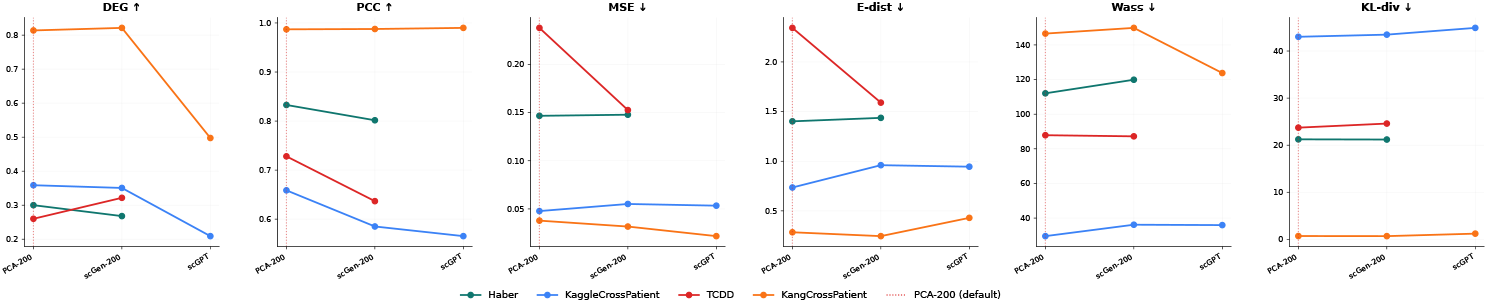
Embedding-method comparison: PCA, scGen, and scGPT. DELPHAI evaluated under three candidate embeddings on the four representative sweep datasets, *o.o.d*. scenario. scGPT is shown only on the two human datasets (KaggleCrossPatient, KangCrossPatient).

**Fig. S3.**
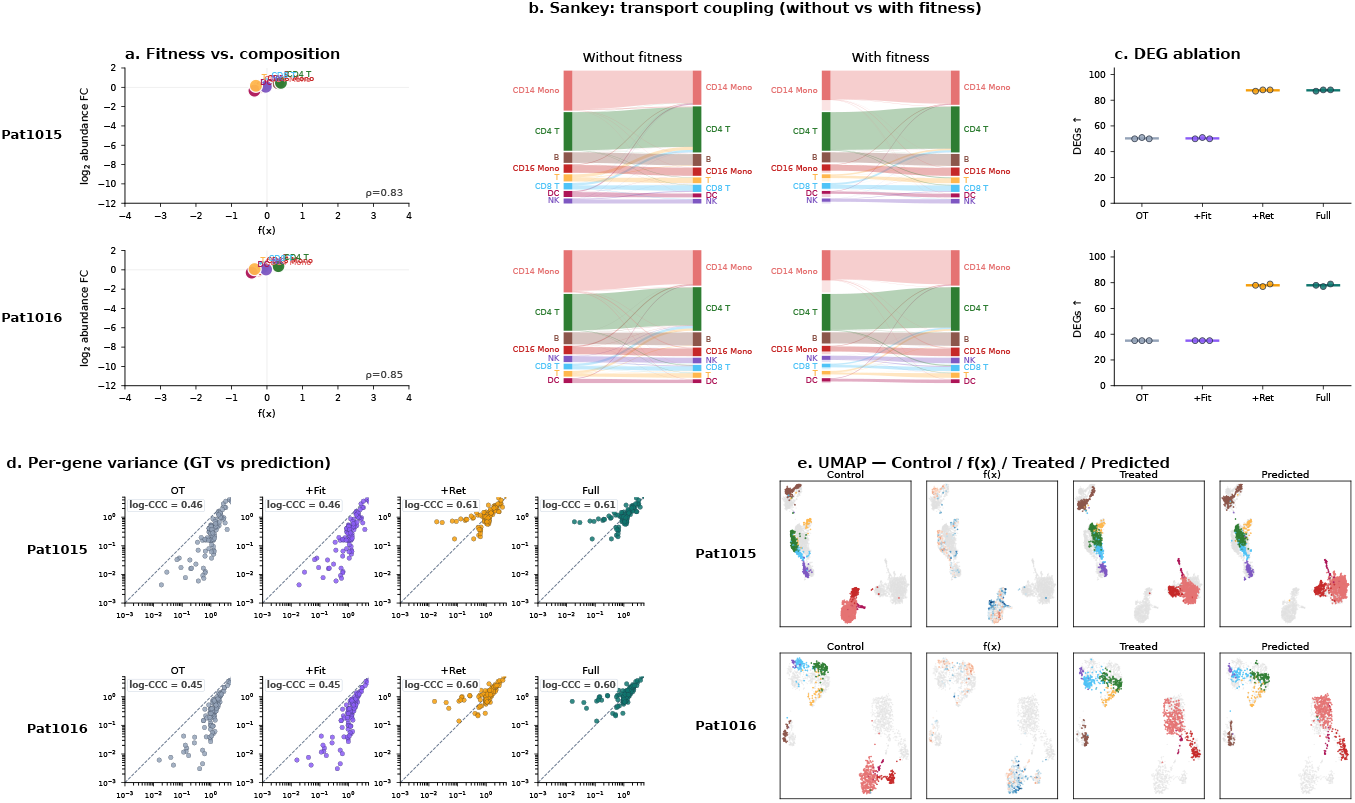
KangCrossPatient (IFN-*β*), additional held-out patients. Same layout as main Fig. 2 (top row), but for held-out patients Pat1015 and Pat1016 from KangCrossPatient.

**Fig. S4.**
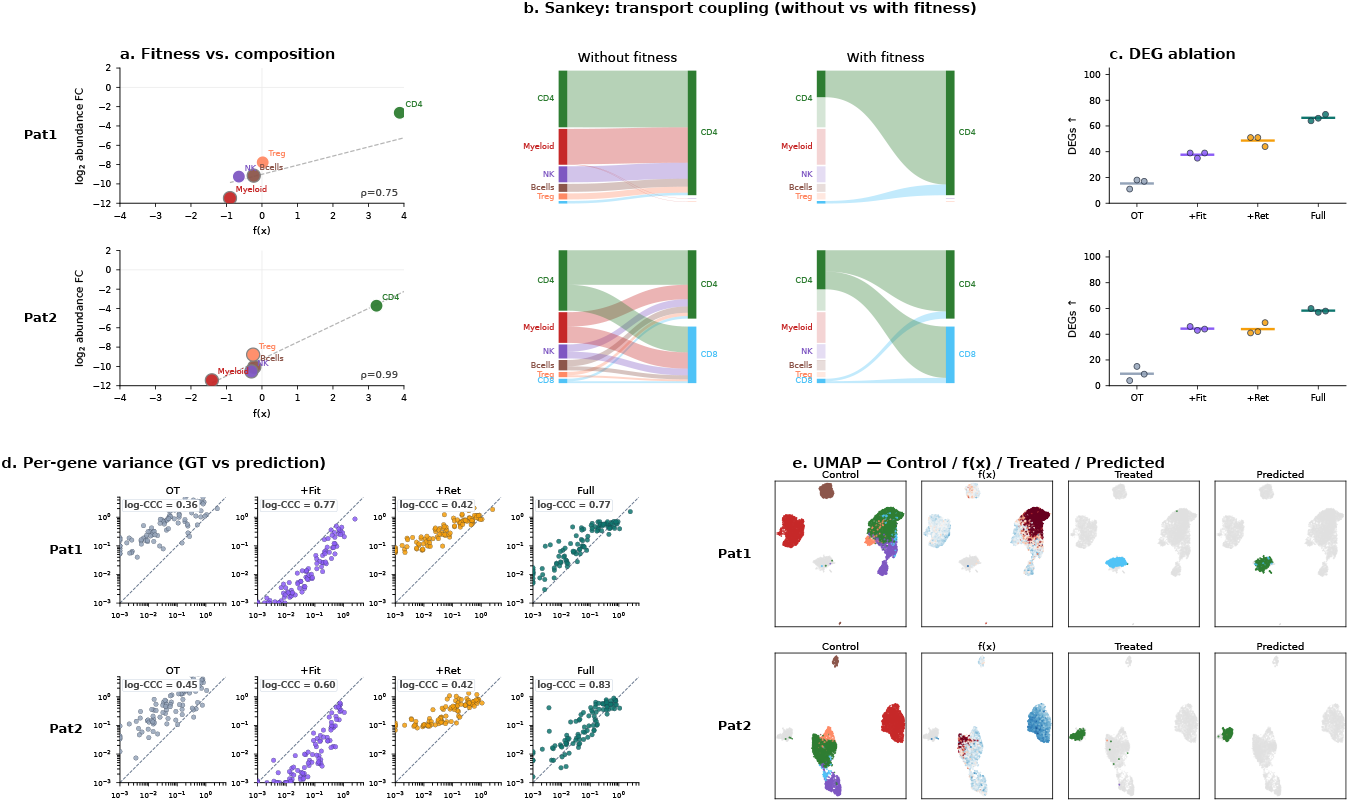
KaggleCrossPatient (CEP18770), additional held-out patients. Same layout as main Fig. 2 (bottom row), but for held-out patients Pat1 and Pat2 from KaggleCrossPatient.

